# A shared pathway of exosome biogenesis operates at plasma and endosome membranes

**DOI:** 10.1101/545228

**Authors:** Francis K. Fordjour, George G. Daaboul, Stephen J. Gould

**Author notes:** **Corresponding author:** Stephen J. Gould, Ph.D., Department of Biological Chemistry, Johns Hopkins University, Baltimore, MD USA.

## Abstract

Eukaryotic cells secrete exosomes, which are small (~30-200 nm dia.), single membrane-bound organelles that transmit signals and molecules to other cells. Exosome-mediated signaling contributes to diverse physiological and disease processes, rendering their biogenesis of high biomedical importance. The prevailing hypothesis is that exosomes bud exclusively at endosome membranes and are released only upon endosome fusion with the plasma membrane. Here we tested this hypothesis by examining the intracellular sorting and exosomal secretion of the exosome cargo proteins CD63, CD9, and CD81. We report here that CD9 and CD81 are both localized to the plasma membrane and bud >5-fold more efficiently than endosome-localized CD63. Furthermore, we show that redirecting CD63 from endosomes to the plasma membrane by mutating its endocytosis signal (CD63/Y235A) increased its exosomal secretion ~6-fold, whereas redirecting CD9 to endosomes by adding an endosome targeting signal (CD9/YEVM) reduced its exosomal secretion ~5-fold. These data demonstrate that the plasma membrane is a major site of exosome biogenesis, and more importantly, that cells possess a common pathway for exosome protein budding that operates at both plasma and endosome membranes. Using a combination of single-particle interferometry reflectance (SPIR) imaging and immunofluorescence (IF) microscopy, we also show that variations in exosome composition are controlled by differential intracellular protein trafficking rather than by separate mechanisms of exosome biogenesis. This new view of exosome biogenesis offers a simple explanation for the pronounced compositional heterogeneity of exosomes and a validated roadmap for exosome engineering.

**Summary:** This study of exosome cargo protein budding reveals that cells use a common pathway for budding exosomes from plasma and endosome membranes, providing a new mechanistic explanation for exosome heterogeneity and a rational roadmap for exosome engineering.

## Introduction

Exosomes are secreted by virtually all eukaryotic cells. These small vesicles are ~30-200 nm dia., have the same topology as the cell, and are enriched in selected proteins, lipids, and nucleic acids(Pegtel and Gould, 2019). Exosomes contribute to numerous physiological processes, including development, immunity, neuronal signaling, etc.(Ashley et al., 2018; Lindenbergh and Stoorvogel, 2018; McGough and Vincent, 2016; Pastuzyn et al., 2018), as well as a wide array human diseases, such as cancer, neurodegeneration, infectious disease, etc.(Becker et al., 2016; Cheng et al., 2018; Gould et al., 2003; van Dongen et al., 2016). Moreover, exosomes are abundant in all biofluids, can be used for clinical liquid biopsies, and are being developed as intrinsic therapeutics as well as drug delivery vehicles(Jia et al., 2014; Li et al., 2017; Phinney and Pittenger, 2017). This breadth of biological importance and translational potential highlights the importance of elucidating the mechanisms of exosome biogenesis.

In their landmark description of secreted vesicles, Trams et al.(Trams et al., 1981) defined exosomes as secreted vesicles that ‘*may serve a physiological purpose*’ and showed that cells secrete two classes of extracellular vesicles (EVs), one that is small, ~40 nm dia., and another that is considerably larger, >500 nm dia. This ~10-fold difference in size allows their separation by differential centrifugation, which led to the eventual restriction of the term *exosome* to mean the smaller class of secreted vesicles, and adoption of the term *microvesicle* to describe the large class of secreted vesicles(Gould and Raposo, 2013). More recently, there has been a concerted effort to redefine these terms yet again, this time on the basis of biogenic mechanism. However, this effort has instead led to redefinition based on the site where an exosome originated in the cell, rather than the actual mechanism of biogenesis, and the now-commonplace dogma that that exosomes arise only by budding into the endosome lumen(Colombo et al., 2014; Crenshaw et al., 2018; Desrochers et al., 2016; Mathieu et al., 2019; van Niel et al., 2018). While there is abundant evidence that exosomes can arise in this manner, the prevailing, ‘endosome-only’ hypothesis of exosomes biogenesis is not supported by clear evidence that exosomes and exosome cargoes cannot bud from the plasma membrane. In fact, it is fair to say that the ‘endosome-only’ hypothesis has yet to be tested.

Highly reductionist, cargo-based studies provided critical early insights into the biogenesis of the endoplasmic reticulum(Blobel and Dobberstein, 1975a; Blobel and Dobberstein, 1975b; Blobel and Sabatini, 1971), nucleus(Kalderon et al., 1984), mitochondria(Horwich et al., 1985), peroxisome(Gould et al., 1989) and other organelles(Blobel, 1980; Blobel, 1995). It is therefore reasonable to use a similar approach in studies of exosome biogenesis. This requires a focus on the most highly enriched exosomal proteins, as these show the strongest evidence of active sorting into exosomes, which are the tetraspanins CD63, CD9, and CD81(Escola et al., 1998; Thery et al., 1999), a trio of proteins that are widely used as exosome markers(Colombo et al., 2014; Crenshaw et al., 2018; Desrochers et al., 2016; Mathieu et al., 2019; van Niel et al., 2018). Like all tetraspanins, these proteins are co-translationally translocated into the ER lumen and membrane, span the membrane 4 times, and have their N-terminus and C-terminus oriented into the cytoplasm. The remainder of this paper tests several predictions of the ‘endosome-only’ hypothesis of exosome biogenesis by following the intracellular sorting and exosomal secretion of these cargo proteins, particularly CD63 and CD9. Our data represent argue against the ‘endosome-only’ model and instead indicate that cells make exosomes by a common mechanism that acts across the spectrum of plasma and endosome membranes.

## Results

A key tenet of the ‘endosome-only’ hypothesis of exosome biogenesis is that exosome cargo proteins must be targeted to the limiting membrane of endosomes as a prerequisite to their exosomal secretion. To assess the validity of this prediction, we first examined the subcellular distribution of three well-established exosome marker proteins, CD63, CD9, and CD81(Escola et al., 1998; Thery et al., 1999). Using immunofluorescence microscopy (IFM), we observed that CD9 and CD81 were highly enriched at the plasma membrane instead of at endosomes, and that only CD63 displayed an endosomal localization (***Fig. 1A***). Although these distributions do not argue against the ‘endosome-only’ hypothesis of exosome biogenesis, they can only be reconciled with this hypothesis if CD63 buds more efficiently into exosomes than either CD9 or CD81. However, when we measured the relative budding of these proteins, we observed the exact opposite result. Specifically, we observed that CD9 displayed a relative budding that was ~5-fold higher than CD63 (4.9 +/- 0.53 fold higher; Student’s t-test *p*-value (*p*) = 0.0053; n = 4) and that CD81 displayed a relative budding ~15-fold higher CD63 (15.7 +/- 2.9 fold higher; *p* = 0.015; n = 4) (***Fig. 1B***).

**Figure 1.**
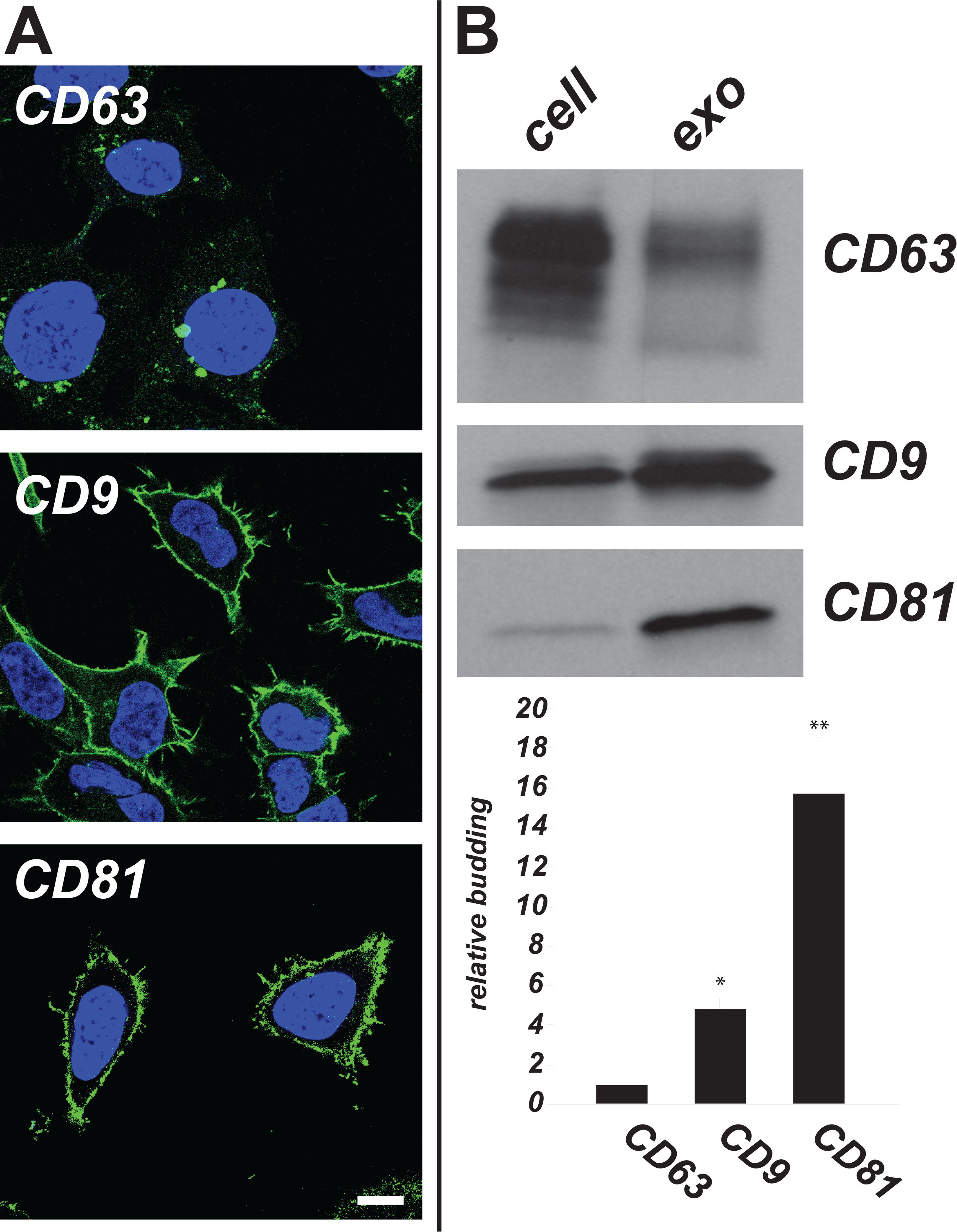
Plasma membrane-localized exosome cargoes CD9 and CD81 display higher relative budding than endosome-localized CD63. (**A**) HEK293 cells were fixed, permeabilized, and stained with antibodies specific for CD63, CD9, or CD81, as well as with DAPI (blue), and then imaged by confocal microscopy. bar, 10 um. (**B**) Immunoblot analysis of HEK293 exosome and cell lysates reveals that the relative budding of CD63 is 5-fold less than that of CD9 and 16-fold less than that of CD81. The bar graph shows the average +/- s.e.m. (n = 4) of the relative budding of CD63, CD9, and CD81 proteins. * denotes a *p*-value less than 0.05, ** denotes a *p*-value less than 0.005.

These observations are inconsistent with the ‘endosome-only’ model of exosome biogenesis and instead raise the possibility that exosome cargo proteins, and thus exosomes, also bud directly from plasma, and that in HEK293 cells they may bud more efficiently from the plasma membrane than from endosome membranes. To differentiate between these models of exosome biogenesis we redirected CD63 from endosomes to the plasma membrane by eliminating it consitiutive endocytosis signal. If the endosome-only hypothesis is correct, then this shift should cause a severe reduction in the exosomal secretion of CD63. However, if HEK293 cells bud exosomes from plasma membranes, the budding of plasma membrane-localized CD63 should remain strong. To execute this experiment, we created plasmids designed to express either WT CD63 or CD63/Y235A, a mutant form of CD63 carrying a mutation that disrupts its clathrin-mediated endocytosis (<u>Y</u>EVM_COOH_, where the Y235 reside critical for binding to the clathrin AP-2 adaptor complex is underlined(Bonifacino and Traub, 2003)). These plasmids were then transfected into CD63^-/-^ cells(Fordjour et al., 2016) and the resulting cells were subsequently processed by IFM to assess the subcellular distribution of WT CD63 and CD63/Y235A. As expected, WT CD63 was localized primarily in endosomes, whereas CD63/Y235A was localized primarily at the plasma membrane (***Fig. 2A***). Exosome and cell lysates were also collected from these cultures, and subsequent IB analysis to assess the relative budding of these proteins. Once again, the data were the exact opposite of that predicted by the endosome-only hypothesis of exosome biogenesis (***Fig. 2B***). Specifically, the plasma membrane-localized CD63/Y235A protein displayed ~6-fold higher relative budding than endosome-localized, WT CD63 (6.1-fold; +/- 1.3-fold, *p* = 0.0038; *n* = 9).

**Figure 2.**
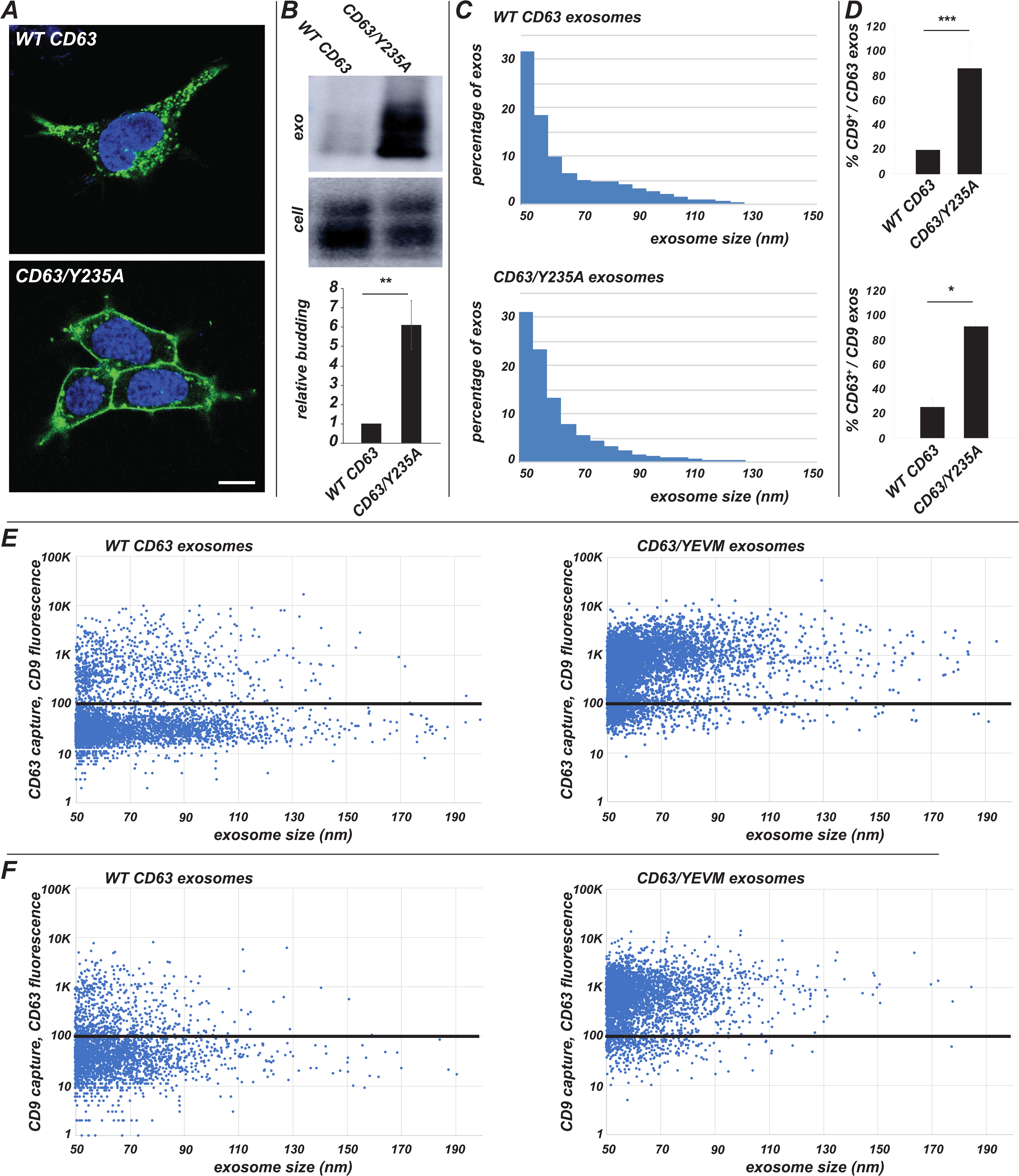
Redirecting CD63 to the plasma membrane enhances its exosomal secretion and its exosomal co-localization with the plasma membrane-enriched exosome marker, CD9. (**A**) Confocal microscopy of HEK293 CD63^-/-^ cells transfected with plasmids expressing either (upper panel) WT CD63 or (lower panel) CD63/Y235A, stained with (green) antibodies specific for CD63 and (blue) DAPI. bar, 10 um. (**B**) Immunoblot analysis of exosome and cell lysates from HEK293 CD63^-/-^ cells expressing either CD63 or CD63/Y235A. Bar graph shows average relative budding +/- s.e.m. (n = 9) of WT CD63 and CD63/Y235A compared to wildtype CD63; *p*-value < 0.005. (**C**) Coupled SPIRI and IFM reveal that (upper graph) WT CD63 exosomes and (lower graph) CD63/Y235A exosomes have similar size distribution profiles. (**D**) Bar graphs showing the percentages (average +/- s.e.m.) of (upper graph) CD9-captured exosomes that stain positive for CD63 and (lower graph) CD63-captured exosomes that stain positive for CD9. Statistical significance is denoted by one asterisk (*p* <0.05), two asterisks (*p* <0.005) or three asterisks (*p* <0.005). (**E**) Scatter analysis plotting CD9 fluorescence (a.u.) vs. exosome size on exosomes captured on anti-CD63 antibodies for (left plot) WT CD63 exosomes and (right plot) CD63/Y235A exosomes. Solid black line denotes the threshold of background fluorescence, as determined by staining samples with non-immune IgG. Black line denotes fluorescence background level of 100. (**F**) Scatter analysis plotting CD63 fluorescence (a.u.) vs. exosome size on exosomes captured on anti-CD9 antibodies for (left plot) WT CD63 exosomes and (right plot) CD63/Y235A exosomes. Black line denotes fluorescence background level of 100.

The most parsimonious interpretation of these data is that cells can make exosomes at both plasma and endosome membranes, and that the plasma membrane is the predominant site of exosome budding in HEK293 cells. If this interpretation is correct, then WT CD63-containing exosomes should be the same size as CD63/Y235A-containing exosomes. To test this prediction, we used single-particle interferometry reflectance (SPIR) imaging(Avci et al., 2015; Daaboul et al., 2017; Daaboul et al., 2016; Sevenler et al., 2017; Sevenler et al., 2018) and immunofluorescence microscopy (IFM) to measure the sizes of thousands of WT CD63 exosomes and CD63/Y235A exosomes. SPIRI-IFM is a novel analytical approach that combines traditional IFM of immunolabeled exosomes with label-free visualization of exosomes using a single-particle interferometric reflectance imaging sensor, in which the interference of light reflected from the sensor surface is (***i***) modified by the presence of a chip-bound exosome, (***ii***) varies in relation to the diameter of the chip-bound exosome, and (***iii***) allows measurement of exosome diameter to a resolution of 0.5 nm. In these experiments, exosomes were collected from the conditioned media of CD63^-/-^ cells expressing either WT CD63 or CD63/Y235A. These were then incubated with SPIRI chips that had been previously functionalized with anti-CD63 antibodies. The chips were washed, the chip surface was incubated with fluorescently labeled antibodies specific for CD63, washed again, and the bound exosomes analyzed by SPIRI-IFM to (a) measure the sizes of thousands of individual exosomes and (b) confirm the exosomal nature of each exosome by the presence of anti-CD63 fluorescence. The resulting data revealed that WT CD63 exosomes and CD63/Y235A exosomes have size distribution profiles that are virtually identical (***Fig. 2C***), and average diameters that are nearly the same (69 nm for WT CD63 exosomes (*n* = 3686) and 65 nm for CD63/Y235A exosomes (*n* = 5569)).

These findings show that cells will bud CD63 in exosomes of the same size, regardless of whether CD63 is localized at endosome membranes or localized at the plasma membrane. The result represents strong evidence that cells possess a common pathway for budding exosomes from both plasma and endosome membranes. If this interpretation is correct, then the composition of individual exosomes will be heavily influenced by the local concentration of exosome cargo molecules. To test this prediction, we asked if WT CD63 exosomes and CD63/Y235A exosomes differed in their inclusion of CD9, a plasma membrane-localized exosome cargo. Specifically, we incubated these two exosome populations on SPIRI chips functionalized with either anti-CD63 or anti-CD9 antibodies, and then probed them with fluorescently-tagged antibodies specific for both CD63 and CD9. The resulting data support this new view of exosome biogenesis as they show pronounced increase in the exosomal co-localization of these proteins on CD63/Y235A-containing exosomes (***Fig. 2D***). For example, when exosomes were captured on anti-CD63 antibodies and interrogated for the presence of CD9, only 20% of WT CD63-containing exosomes stained positive for CD9 (20% +/- 1%) whereas >80% of CD63/Y235A exosomes stained positive for CD9 (86% +/- 21%), a 4.3-fold increase (*p* = 0.00011; n = 3). Similar results were observed when these exosomes were captured on anti-CD9 antibodies and stained for CD63, which revealed an exosomal co-localization of CD63 on a quarter (25% +/- 7%) of CD9-captured exosomes from WT CD63-expressing cells, and an exosomal co-localization of CD63 on >90% of CD9-captured exosomes from cells expressing CD63/Y235 (91% +/- 5%), a 3.6-fold increase (*p* = 0.032; n = 3).

The pronounced increase in the exosomal co-localization of CD9 on CD63/Y235A exosomes vs WT CD63 exosomes can also be visualized by plotting fluorescence intensity for thousands of individual exosomes (***Fig. 2E, F***). This is evident from comparing plots of anti-CD9 fluorescence intensity for (left panel) WT CD63 exosomes captured on anti-CD63 antibodies to those of (right panel) CD63/Y235A exosomes captured on anti-CD63 antibodies (***Fig. 2E***). It can also be seen in plots of anti-CD63 fluorescence intensity for (left panel) WT CD63 exosomes captured on anti-CD9 antibodies and (right panel) CD63/Y235A exosomes captured on anti-CD9 antibodies (***Fig. 2F***).

The simplest interpretation of the above data is that CD63 and CD9 bud by a common mechanism that operates across the spectrum of endosome and plasma membranes, and that the composition of any individual exosome is determined in large part by the local concentration of other exosome cargoes. If this model is correct, then the same trends should be observed upon redirecting CD9 from plasma to endosome membrane. Towards this end, we generated CD9^-/-^ HEK293 cells (***supplementary figure S1***) and then transfected them with with plasmids designed to express either WT CD9 or CD9/YEVM, a form of CD9 that carries the constitutive endocytosis signal from CD63 (<u>Y</u>EVM_COOH_). Immunofluorescence microscopy of these cells confirmed that WT CD9 was correctly localized to the plasma membrane and that CD9/YEVM was redirected to endosome membranes (***Fig. 3A***). Furthermore, IB analysis of cell and exosome fractions prepared from these cell populations revealed that redirecting CD9 from the plasma membrane to endosomes resulted in an ~5-fold reduction in its relative budding compared to that of WT CD9 (***Fig. 3B***; 5.5-fold difference; *p* = 0.00019; n = 9), demonstrating once again that the plasma membrane-localized form of a protein buds better than its endosome-targeted counterpart.

**Figure 3.**
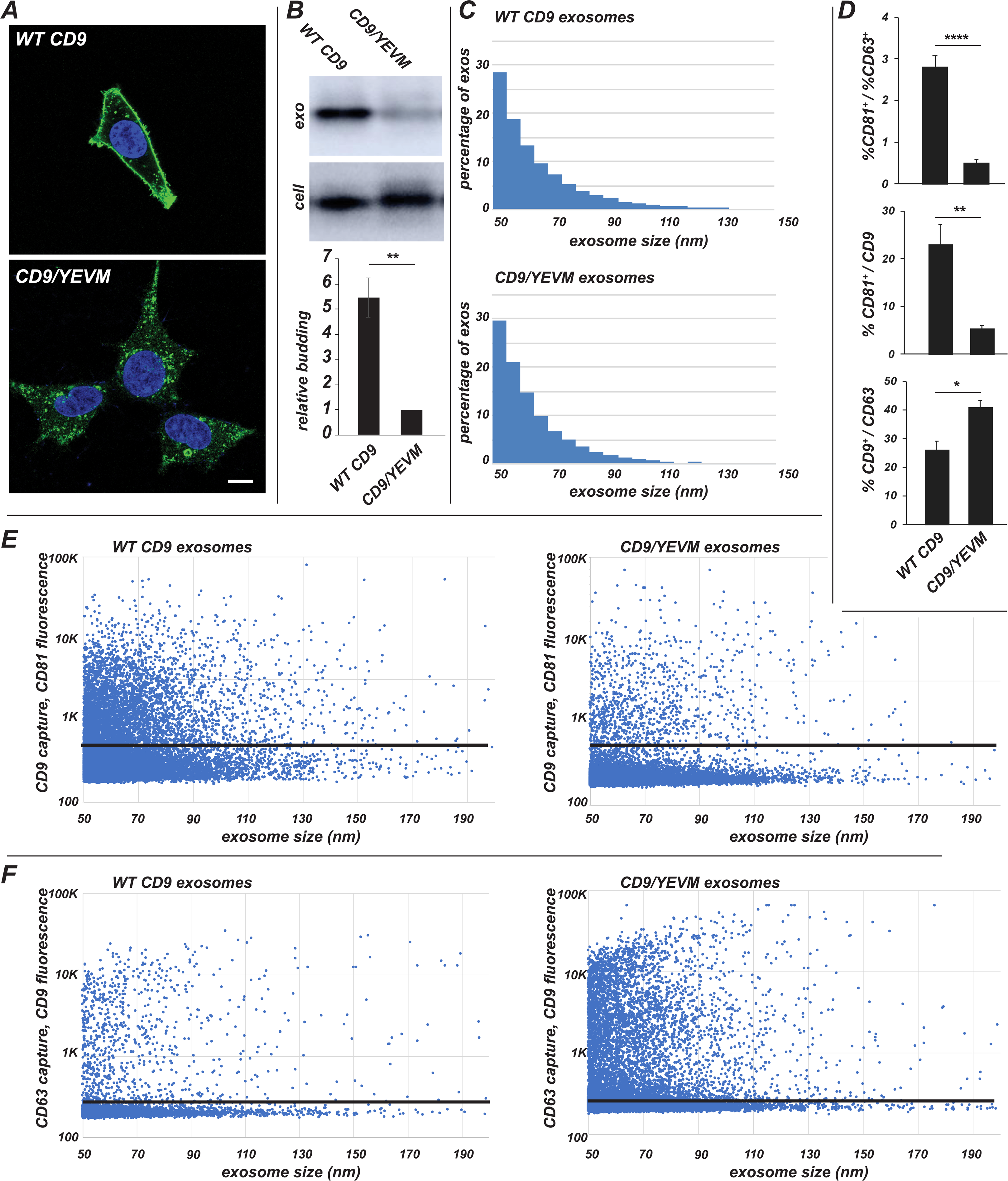
Redirecting CD9 to endosomes inhibits its exosomal secretion, reduces its exosomal co-localization with CD81, and increases its exosomal co-localization with CD63. (**A**) Confocal microscopy of HEK293 CD9^-/-^ cells transfected with plasmids expressing either (upper panel) wildtype CD9 or (right panel) CD9/YEVM, stained with (green) antibodies specific for CD9 and (blue) DAPI. bar, 10 um. (**B**) Immunoblot analysis of cell and exosome fractions from HEK293 CD9^-/-^ cells expressing either CD9 or CD9/YEVM. Graph shows average relative budding +/- s.e.m. (n = 9) of CD9 compared to CD9/YEVM; *p*-value <0.0005. (**C**) Size distribution of (upper graph) CD9-containing exosomes (upper graph) and (lower graph) CD9/YEVM-containing exosomes, as determined by coupled SPIRI and IFM. (**D**) Bar graphs showing (upper graph) the ratio of CD81-positive exosomes to CD63-positive exosomes captured by anti-CD9 antibodies from CD9 exosomes and CD9/YEVM exosomes; (middle graph) the percentage of CD81-positive exosomes captured on anti-CD9 antibodies from CD9 exosomes and CD9/YEVM exosomes; and (lower graph) the percentage of CD9-positive exosomes captured on anti-CD63 antibodies from CD9 exosomes and CD9/YEVM exosomes. Each value represents that average +/- s.e.m., and the degree of statistical significance is denoted by one asterisk (*p* <0.05), two asterisks (*p* <0.005) or four asterisks (*p* <0.0005). (**E**) Scatter analysis of CD81 fluorescence intensity (a.u.) plotted against exosome diameter, of exosomes captured on anti-CD9 antibodies from (left plot) WT CD9 exosomes and (right plot) CD9/YEVM exosomes. Black line denotes fluorescence background level of 300. (**F**) Scatter analysis of CD9 fluorescence intensity (a.u.) plotted against exosome diameter, of exosomes captured on anti-CD63 antibodies from (left plot) WT CD9 exosomes and (right plot) CD9/YEVM exosomes. Black line denotes fluorescence background level of 250.

To further characterize WT CD9 exosomes and CD9/YEVM exosomes, we once again employed SPIRI-IFM. WT CD9 exosomes and CD9/YEVM exosomes were incubated with to anti-CD9 antibody-functionalized SPIRI chips, bound exosomes were then stained with anti-CD9 antibodies, and the chips were interrogated by SPIRI-IFM. The resulting data revealed that WT CD9 exosomes and CD9/YEVM exosomes have similar size distribution profiles (***Fig. 3C***) and average diameters (67 nm for WT CD9 exosomes (n = 15,684) and 65 nm for CD9/YEVM exosomes (n = 18,232)), demonstrating that these proteins bud from the cell in bona fide exosomes.

The budding of both CD9 and CD9/YEVM in exosomes of the same size supports the idea that cells make exosomes from both plasma and endosome membranes. If this interpretation is correct, then the antigenic character of WT CD9 exosomes and CD9/YEVM exosomes should differ in respect to CD81 and CD63, with WT CD9 exosomes showing a higher degree of exosomal co-localization with plasma membrane-enriched CD81, and CD9/YEVM showing a higher degree of exosomal co-localization with endosome-localized CD63. To test this prediction, WT CD9 exosomes and CD9/YEVM exosomes were bound to anti-CD9-functionalized SPIRI chips, incubated with antibodies specific for CD81 and CD63, and subjected to SPIRI-IFM (***Fig. 3D***). The resulting data revealed that the ratio of CD81-positive exosomes to CD63-positive exosomes was 2.8 (+/- 0.3) for WT CD9 exosomes, but fell 5.5-fold for CD9/YEVM exosomes to 0.51 (+/- 0.06), a decrease of high significance (*p* = 0.000020; n = 9). This shift in antigenic character is also reflected in a 4-fold decrease (*p* = 0.0037; n = 9) in the percentage of CD9-containing exosomes that contain CD81 (from 23% +/- 4% for WT CD9 exosomes to 5.4% +/- 0.7% for CD9/YEVM exosomes). It is also evident in a 1.7-fold increase (p = 0.011, n = 9) in the percentage of CD63-captured exosomes that contain CD9 (from 26% +/- 3% for WT CD9 exosomes to 41% +/- 2% for CD9/YEVM exosomes).

These changes in the exosomal co-localization of CD81, CD63 and CD9 can also be visualized by scatter plots of fluorescence intensity for thousands of individual exosomes (***Fig. 3E, F***). This is particularly evident from plotting anti-CD81 fluorescence intensity (a.u.) for (left panel) WT CD9-exosomes captured on anti-CD9 antibodies and (right panel) CD9/YEVM exosome captured on anti-CD9 antibodies (***Fig. 3E***). It can also be seen by plotting anti-CD9 fluorescence intensity (a.u.) for (left panel) WT CD9 exosomes captured on anti-CD63 antibodies and (right panel) CD9/YEVM exosomes captured on anti-CD63 antibodies (***Fig. 3F***).

Our model is agnostic on the point of whether cells bud exosomes predominantly from the plasma or endosome membranes, as we presume that both are possible. Nevertheless, the observation that HEK293 cells display such a pronounced preference for making exosomes from the plasma membrane raises the question of whether this is also true for any other cell types. To answer this question we expressed the WT CD63, CD63/Y235A, WT CD9, and CD9/YEVM proteins in mouse NIH3T3 fibroblasts and followed their intracellular sorting and exosomal secretion (***Fig. 4***). More specifically, we transfected NIH3T3 cells with plasmids designed to express each protein, selected for G418-resistant clones, and assayed each for levels of expression to obtain matched pairs of cell lines that express similar levels of each cargo protein. Each cell line was then subjected to IFM to determine the subcellular distribution of the protein of interest, and immunoblot analysis was used assess their relative budding. As expected, CD63/Y235A and WT CD9 were enriched at the plasma membrane whereas WT CD63 and CD9/YEVM were targeted to endosomes (***Fig. 4A-D***). Furthermore, we observed that redirecting CD63 from endosomes to the plasma membrane of NIH3T3 cells led to a significant increase in its relative budding from the cell (5.7 +/- 0.7 fold, *p* = 0.00058, n = 7; ***Fig. 4E***), similar to what we observed for HEK293 cells. The same was true for the CD9 experiments, as and redirecting CD9 from the plasma membrane to endosomes of NIH3T3 cells once again led to a significant decrease in its relative budding from the cell (3.3 +/- 0.6 fold, *p* = 0.0056, n = 8; ***Fig. 4F***). Thus, NIH3T3 resemble HEK293 cells in their preferential exosomal secretion of plasma membrane-targeted exosome cargoes.

**Figure 4:**
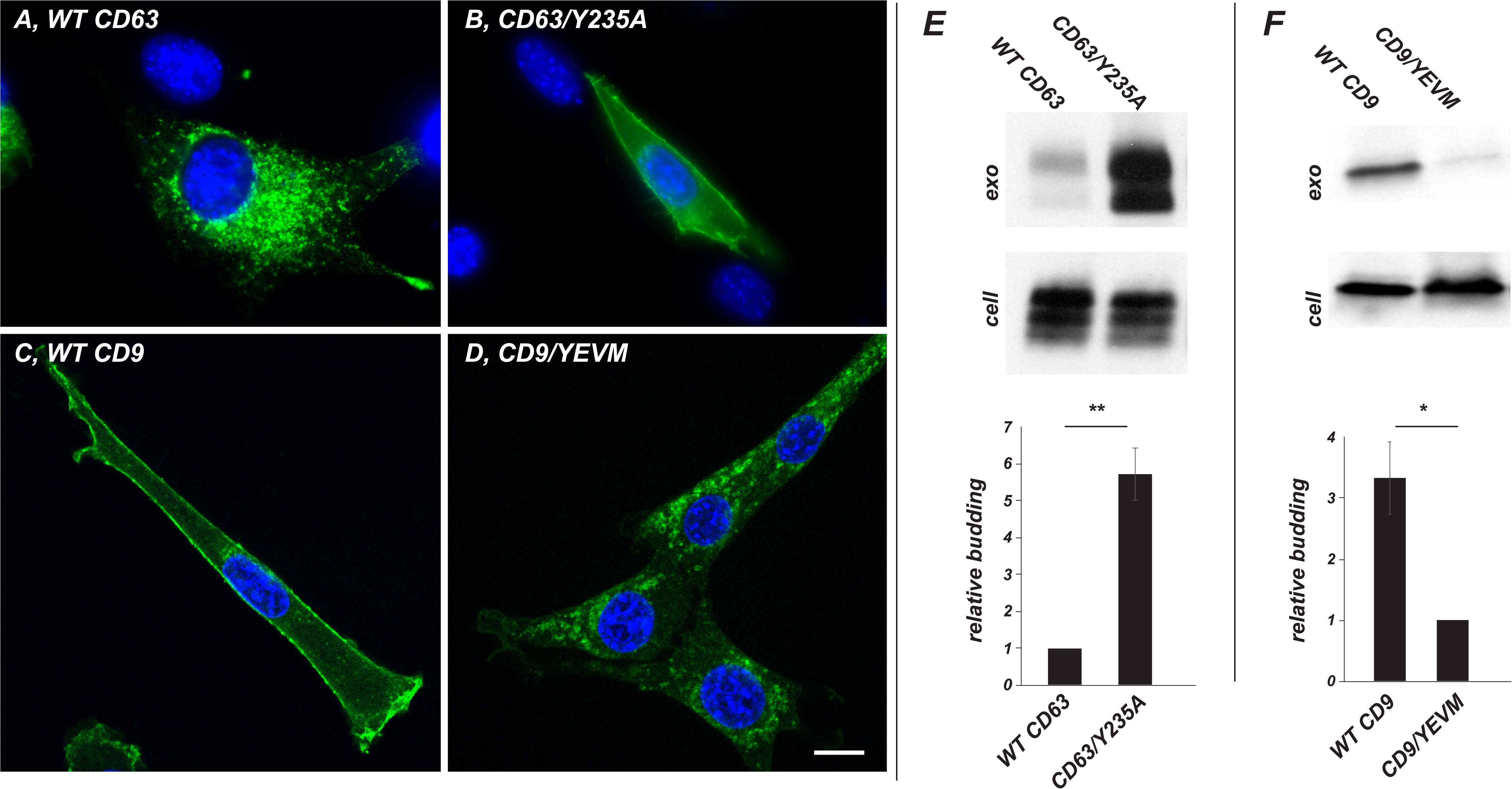
NIH3T3 cells bud exosome cargoes preferentially from plasma membranes. (**A-D**) IFM of NIH3T3 cells stably expressing (A) WT human CD63, (B) human CD63/Y235A, (C) WT human CD9, or (D) human CD9/YEVM. Cells were fixed, permeabilized, incubated with monoclonal antibodies specific for (A,B) human CD63 or (C,D) human CD9, stained with secondary antibodies specific for mouse IgG and also with DAPI to detect the nucleus. Bar, 10 μm. (**E**) Immunoblot analysis of cell and exosome fractions collected from NIH3T3 cells stably expressing WT CD63 or CD63/Y235A, followed by calculation of relative budding and appropriate statistical analysis. (**F**) Immunoblot analysis of cell and exosome fractions collected from NIH3T3 cells stably expressing WT CD9 or CD9/YEVM, followed by calculation of relative budding and statistical analysis.

## Discussion

### Testing the ‘endosome-only’ hypothesis

A wide array of reviews and research articles assert that exosomes arise solely by budding into endosomes, followed by endosome-plasma membrane fusion(Colombo et al., 2014; Crenshaw et al., 2018; Desrochers et al., 2016; Mathieu et al., 2019; Thery et al., 2018; van Niel et al., 2018). We find this perplexing, in part because there is no compelling body of evidence that exosomes cannot bud from the plasma membrane, and in part because no study has even attempted to test this prevailing, ‘endosome-only’ hypothesis. Here we subjected this hypothesis to experimental interrogation, primarily by testing its core tenet that exosomal cargoes must be targeted to endosomes before they can be secreted in exosomes.

The foundation of the ‘endosome-only’ hypothesis is belied by the fact that CD9 and CD81, which display the highest relative budding of any exosomal cargo proteins yet reported, are localized primarily at the plasma membrane, whereas endosome-targeted CD63 buds 5-15-fold less efficiently that either CD9 or CD81. After all, if exosomes only bud via endosomal budding, then cargoes that are enriched at endosomes should bud more efficiently than plasma membrane-localized cargoes. This tenet is also contradicted by the consequences of redirecting CD63 to the plasma membrane. Under the prevailing paradigm, this change in subcellular distribution should reduce or eliminate the exosomal secretion of CD63, while in reality it led to a ~6-fold increase in its exosomal secretion. The prevailing paradigm was similarly unable to predict or explain the consequence of redirecting CD9 from the plasma membrane to endosome membranes, which resulted in an ~5-reduction in CD9’s exosomal secretion. In fact, every observation in this paper runs counter to the assertion that exosomes are generate exclusively by budding from the endosome membrane. Given its consistent inability to predict the outcome of simple empirical tests, one cannot help but wonder whether the prevailing paradigm is based on anything more than a circular argument in which exosomes are believed to arise by endosomal budding for the sole reason that exosomes have been defined in that manner.

### Exosomes arise from plasma and endosome membranes

The simplest interpretation of our data is that cells possess a shared mechanism of exosome biogenesis that operates at plasma and endosome membranes, and that HEK293 and NIH3T3 cells make most of their exosomes at the plasma membrane. This hypothesis predicts or explains all major observations in this report, including:

i. the steady-state enrichment of exosome cargoes at both plasma and endosome membranes;
ii. the ~5-15-fold higher relative budding of CD9 and CD81 relative to CD63;
iii. the exosomal secretion of CD63/Y235A;
iv. the ~6-fold higher exosomal secretion of CD63/Y235A compared to WT CD63;
v. the ~5-fold lower exosomal secretion of CD9/YEVM compared to WT CD9;
vi. the exosomal co-localization of CD9 and CD63 on WT CD63 exosomes;
vii. the ~4-fold increase in exosomal co-localization of CD9 and CD63 on CD63/Y235A exosomes compared to WT CD63 exosomes;
viii. the ~4-fold decrease in exosomal co-localization of CD81 on CD9/YEVM exosomes compared to WT CD9 exosomes; and
ix. the ~1.7-fold increase in CD63 on CD9/YEVM exosomes compared to WT CD9 exosomes.

The model of exosome biogenesis posited here is also consistent with prior observations of exosome biogenesis via the plasma membrane(Anderson, 1969; Anderson et al., 2005; Bianchi et al., 2014; Booth et al., 2006; Cantaluppi et al., 2012; Casado et al., 2017; Fang et al., 2007; Shen et al., 2011a; Shen et al., 2011b) as well as prior observations of exosome biogenesis via endosome membranes(Colombo et al., 2014; Crenshaw et al., 2018; Desrochers et al., 2016; Harding et al., 1983; Harding et al., 1984; Mathew et al., 1995; Mathieu et al., 2019; Pan and Johnstone, 1983; Pegtel and Gould, 2019; van Niel et al., 2018). Furthermore, it adheres to the principle of maximum parsimony by offering a simpler explanation for a wider array of data than is possible under the ‘endosome-only’ hypothesis of exosome biogenesis.

### Implications for exosome composition and heterogeneity

Although our data indicate that a single mechanism of exosome biogenesis operates across the spectrum of endosome and plasma membranes, this model does not predict a uniform composition of secreted exosomes. To the contrary, it predicts that the composition of each individual exosome will be determined primarily by the local, nanometer-scale concentrations of exosome cargoes in the immediate vicinity of each nascent exosome, which are then fixed by its scission from its parent membrane. This prediction is supported by several of our observations, including the increased exosomal co-localization of CD9 on CD63/Y235A exosomes, the decreased exosomal co-localization of CD81 on CD9/YEVM exosomes, and the increased exosomal co-localization of CD63 on CD9/YEVM exosomes.

This model also provides a simple yet elegant explanation for the pronounced compositional heterogeneity of exosomes as the vesicular manifestation of nanometer-scale heterogeneities in the plasma and endosome membranes that give rise to exosomes. Compositional heterogeneity of these membranes is an established fact(Bernardino de la Serna et al., 2016; Sevcsik et al., 2015; Sevcsik and Schutz, 2016; Specht et al., 2017), and these heterogeneities are generated by both mechanistic and stochastic forces. For example, the protein sorting machineries that distribute cargoes along the spectrum of endosome and plasma membranes display significant-to-subtle differences in affinity for virtually every protein with which they interact(Traub and Bonifacino, 2013), resulting a spectrum of large-scale distribution patterns of these proteins along the spectrum of plasma and endosome membranes. In addition, each cargo molecule will experience a range of stochastic forces further affecting their nanometer-scale concentration at these membranes, including diffusion, structural fluctuations, and various intermolecular interactions. The combination of these forces, together with the varied size of exosomes, indicates that each exosome cargo molecule will emerge at various levels on exosomes of varied size, with few if any having the exact same size and amount of any one protein. This prediction corresponds rather well with our SPIRI-IFM data, in which we interrogated thousands of HEK293-derived exosomes for just two parameters, size and CD9 abundance, and found that each exosome had a nearly unique combination of size and CD9 staining. If one assumes that a similar heterogeneity is seen for the thousands of proteins, nucleic acids, and lipids that are incorporated into exosomes by a single cell line(Li et al., 2016), it may be that no two exosomes are alike.

It should be noted that this model predicts additional contributors to structural and functional heterogeneity. For example, exosomes that arise by budding from the plasma membrane are likely influenced by the high complexity of the extracellular milieu, whereas those that bud into endosomes are affected by its low pH and abundance of hydrolytic products (e.g. peptides, lipids, etc.). Exosomes may also arise by budding into intracellular plasma membrane-connected compartments (IPMCs), and this environment may also affect exosome composition and function(Nkwe et al., 2016; Pelchen-Matthews et al., 2012). IPMCs are virtually indistinguishable from multivesicular bodies (MVBs) but are not endosomes at all, as their lumens are contiguous with the extracellular space and their membranes are continuous with the cell surface. These different sites of origin can also impart a temporal heterogeneity in exosome release, as vesicles retained within MVBs or IPMCs can be released in a delayed and pulsatile manner via endosome fusion with the plasma membrane(Sung et al., 2015; Verweij et al., 2018) and IPMC opening(Pelchen-Matthews et al., 2012), respectively.

### Applications to exosome engineering

The mechanisms of exosome biogenesis have important implications for the more prosaic topic of exosome engineering. Exosomes are being developed as therapeutics and drug-delivery vehicles for a number of applications(Gilligan and Dwyer, 2017; Kamerkar et al., 2017; Li et al., 2017), some of which involve genetic engineering of exosome-producing cells(Yim et al., 2016). Efficient exosome engineering requires a solid understanding of exosome biogenesis, as the pathway of exosome biogenesis is the blueprint for exosome design. From this perspective, the ‘endosome-only’ hypothesis of exosome biogenesis does not appear to be particularly helpful, as it wrongly predicts that putative new exosome cargoes should be sent to the endosome membrane. In contrast, the hypothesis posited here provides a validated roadmap for the first two steps in exosome engineering. Specifically, it suggests that exosome engineers (i) test whether the exosome producing cell line of choice makes exosomes predominantly from plasma or endosome membranes, and then (ii) engineers the putative new exosomal cargo so that cells direct it too that location. However, additional steps are also likely involved, in part because most plasma membrane proteins are not targeted to exosomes(Escola et al., 1998; Thery et al., 1999) and in part because protein targeting to exosomes is induced by high-order oligomerization of plasma membrane-anchored proteins(Fang et al., 2007).

### Two-pathway alternatives

It is also useful to consider whether a two-mechanism model could explain the biogenesis of exosomes. For example, is it possible that cells possess one pathway that mediates the exosomal secretion of ‘class A’ cargoes, such as CD9 and CD81, and a second, separate pathway that mediates the budding of ‘class B’ cargoes, such as CD63? To answer this question it is first necessary to accept that the word *separate* means distinct and non-overlapping. This is obvious from its dictionary definition, but also from the cell biological principle of protein topogenesis(Blobel, 1980; Blobel, 1995), in which separate organelles are defined in large part by their mechanistic connection to separate protein sorting pathways. This means that two-pathway models of exosome biogenesis demands that the exosomal co-localization of Class A and Class B cargoes is either minimal or absent.

The data in our paper argue strongly against a two-pathway model of exosome biogenesis, for the simple reason that we detected significant exosomal co-localization of Class A and Class B cargoes in every exosome sample we examined. For example, exosomal co-localization of CD9 and CD63 was observed on a relatively low but nonetheless highly significant percentage of exosomes, as ~25% of WT CD63 exosomes contained both of these Class A and Class B cargoes. Furthermore, their exosomal co-localization could be increased further, rising to ~90% merely by redirecting CD63 to the plasma membrane. Significant exosomal co-localization of Class A and B cargoes was also evident on a significant percentage (~20%) of WT CD9 exosomes, which was doubled to ~40% by redirecting CD9 to endosomes.

Another possible permutation of the two-pathway model would be to restate the Class A pathway as mediates the budding of proteins from the plasma membrane while a Class B pathway mediates protein budding from endosomes. This version of the two-pathway model posits that misdirecting plasma membrane cargoes to the endosome should inhibit its exosomal secretion, which actually matches the data we obtained by redirecting CD9 to endosomes (a 5-fold reduction in CD9 budding). However, this hypothesis also predicts that CD63 budding should also decrease upon its redistribution to the plasma membrane, which runs counter to the fact that it induced a ~6-fold increase in its exosomal secretion. It also predicts a dearth of exosomal co-localization of CD9 and CD63, and is similarly undermined by the contradictory observations outlined in the preceding paragraph.

In light of these considerations it seems that the only way to square our data with a two-mechanism model of exosome biogenesis would be to: (i) reject the conventional meaning of the word ‘separate’ and redefine it to mean ‘pretty much the same’; (ii) ignore the well-established principles of protein topogenesis, in which organelle identity is driven by non-overlapping protein sorting pathways(Blobel, 1980; Blobel, 1995); and (iii) posit that cells somehow evolved two separate pathways of exosome biogenesis that accept the same sets of cargo proteins, secrete them from the cell in exosomes of the same size and molecular characteristics, and have no discernable difference. It seems far more logical to adhere to the principle of maximum parsimony and go with the far simpler model, in which cells use a shared pathway for exosome biogenesis from plasma and endosome membranes.

## Acknowledgments

We thank Colin Fowler, James Morrell, and Jerry Plange of the Gould lab and Aditya Dhande of Nanoview Sciences for expert technical assistance. This work was supported by NIH (U19CA179563).

## Author contributions

F.K.F. performed all experiments other than the coupled SPIRI-IFM analyses, contributed to the experimental design, data interpretation and writing of the paper; G.G.D. performed all SPIRI-IFM analyses and contributed to the experimental design, data interpretation and writing of the paper; S.J.G. conceived of the project, contributed to the experimental design, data interpretation, performed the majority of the SPIRI-IFM analysis, and composition of the manuscript.

## Methods

### Plasmids

The CD9 knock-out plasmid pJM1084 was generated by inserting CD9 gene-specific guide RNA-encoding sequences specific for CD9 exons 1 and 3, respectively, were inserted downstream of the 7sk and H1 promoters, respectively, of pFF4(Fordjour et al., 2016). Plasmids pCF1, pCF2, pCF6, and pCF7 were created by inserting the ORFs encoding CD63, CD63/Y235A, CD9, and CD9/YEVM downstream of the CMV promoter of pcDNA3. Plasmids were amplified by growth in *E. coli* DH10 and purified by ion exchange chromatography from bacterial cell lysates.

### Cells, transfections

HEK293 cells (ATCC CRL1573), HEK293 CD63^-/-^ cells(Fordjour et al., 2016), HEK293 CD9^-/-^ cells and NIH3T3 cells (ATCC CRL-1658) were grown in DMEM supplemented with 10% fetal calf serum (FCS) or 10% exosome-free FCS, with all exosome-related studies performed using cells grown in the latter. HEK293 cells carrying null mutations in CD9 (CD9^-/-^) were generated by transfecting HEK293 cells with pJM1084, selecting for puromycin-resistant cells (7 days), followed by limiting cell dilution into 96 well plates and expansion of single cell clones (SCCs). Multiple independent SCCs were screened by IB using anti-CD9 antibodies, followed by genomic DNA extraction, PCR analysis using oligos that flank both sides to the exon 1 and exon 3 target sites, and sequence analysis of individual PCR products. The CD9^-/-^ cell line selected for further analysis here was found to carry only null alleles in the CD9 gene. Transfections were performed using lipofectamine 2000 (ThermoFisher) according to the manufacturer’s instruction. NIH3T3 cells expressing WT CD63, CD63/Y235A, WT CD9 or CD9/YEVM were selected in 400 μg/ml G418 (ThermoFisher), clones expressing similar level of each pair protein were identified, and then used for subsequent experiments.

### Exosome purification and immunoblot

For each trial, 6 x 10^6^ cells were seeded onto 2 x 150 mm dishes in a total volume of 60 ml of DMEM supplemented with 10% exosome-free FCS and grown for 72 hrs. For all exosome studies, the tissue culture media was spun at 5000 x *g* for 15 min. The pellet was discarded and the supernatants (SN) were passed through 0.22 um filter. For exosome analysis by SPIRI and IFM, the filtrate was concentrated by angular flow filtration (Centricon Plus-70; EMDMillipore) to a final volume of to 500ul. Exosomes were purified by size exclusion chromatography (Izon qEV column), 500ul fraction samples were collected, and fractions 4, 5, and 6 were assayed by immunoblot (IB) to confirm the presence of exosome markers, pooled, and interrogated by SPIRI and IFM. For exosome analysis by IB, the clarified tissue culture supernatant was spun twice at 10,000 x *g* for 30 mins to remove contaminating microvesicles, and the resulting supernatant was spun at 70,000 x *g* for 2 hrs at 4°C to pellet exosomes. Cell lysates were generated by addition of 2 ml of 2x SDS-PAGE sample buffer. Exosome pellets were resuspended in 600ul of 2x SDS-PAGE sample buffer. Immunoblots were performed at a constant ratio of exosome:cell lysates. IB analysis was performed by separating cell and exosome lysates by SDS-PAGE. Proteins were then transferred to Immobilon membranes (EMDMillipore), followed in sequence by incubation with block solution (0.2% non-fat dry milk in TBST), primary antibody solution, 5 washes with TBST, secondary antibody solution, and 5 washes with TBST. Antigens were visualized by chemiluminescence and detected using an Amersham Imager 600 (GE Healthcare Life Sciences) gel imaging system. The resulting digitized IB images were then processed in Image J by converting them to 8-bit grayscale files followed by background subtraction. Measurement parameter and scale were set to integrated density and pixel, respectively. Images were then inverted, bands were delineated using the freehand selection tool, and signal densities were converted to relative protein abundance by multiplying by the dilution factor for each sample. Relative budding was calculated by dividing the protein abundance in exosome lysate by the sum of the protein abundance in the cell lysate and the protein abundance in exosome lysate.

### Immunofluorescence microscopy

Immunofluorescence microscopy (IFM) was performed on cells grown on cover glasses. Cells were fixed (3.7% formaldehyde in PBS for 15 min.), permeabilized (1% Triton X-100 in PBS for 5 min.), incubated with primary antibodies in PBS (15 min.), washed 3 times with PBS, incubated with fluorescently-labeled secondary antibody and DAPI, washed 3 times with PBS, mounted on glass slides, and visualized by confocal microscopy. Antibodies were diluted in PBS (1:200 dilution for CD63 (clone E-12, #sc-365604, Santa Cruz Biotechnology), 1:200 dilution for CD9 (clone H19a, #312102, Biolegend), 1:1000 dilution of fluorescein (FITC) AffiniPure Goat Anti-Mouse IgG (H+L) (#115-095-003 Jackson Laboratory). Confocal images were acquired using a Zeiss AxioObserver inverted microscope with LSM700 confocal module and 63x, 1.4 aperture AxioPlan objective. Images were acquired using Zen software, converted to tiff files, and imported into Adobe Photoshop and Illustrator to create final images. Standard immunofluorescence imaging of NIH3T3 cells was performed at room temperature on a BH2-RFCA microscope (Olympus) equipped with an Olympus S-plan Apo 63× 0.40 oil objective and a Sensicam QE (Cooke) digital camera using IPLab 3.6.3 software (Scanalytics, Inc.). Again, tiff images were imported into Adobe Photoshop and Illustrator to create final images.

### SPIRI and IFM analysis

Each exosome sample was diluted ten-fold in SPIRI incubation buffer (50mM HEPES, 150 mM NaCl and 0.05% Tween-20, pH 7.3). 35 µL of each sample were then incubated on the ExoView Tetraspanin Chip (EV-TC-TTS-01) placed in a sealed 24 well plate for 16 hours at room temperature. Each chip was then washed on an orbital shaker once with PBST (PBS supplemented with 0.05% Tween-20) for 3 minutes, then washed three additional times with PBS for 3 minutes each Chips were then incubated with one or more of Alexa-55-labeled anti-CD81, Alexa-488-labeled anti-CD63, and Alexa-647anti-CD9 antibodies in PBST supplemented with 2% BSA in a volume of 250 µL for 2 hours at room temperature without shaking. Each chip was then washed once with PBST, 3 times with PBS, once in filtered deionized water, and then dried at room temperature for 1 hour. The chips were then imaged with the ExoView R100 reader using the ExoScan 2.5.5 acquisition software (Nanoview Biosciences). The resulting size and fluorescence intensity information for each individual exosome was exported to Excel for statistical analyses. Fluorescence values are reported in arbitrary units.

### Data analysis and presentation

All quantitative data is reported as average +/- standard error of the mean. The statistical significance of differences between different data sets was assessed using Student’s t-test (two-tailed, paired). Histograms and scatter plots were generated using Excel. Images were imported into Adobe photoshop and figures were assembled in Adobe Illustrator. Image data was adjusted for brightness only.

---

**Supplementary Figure 1:**
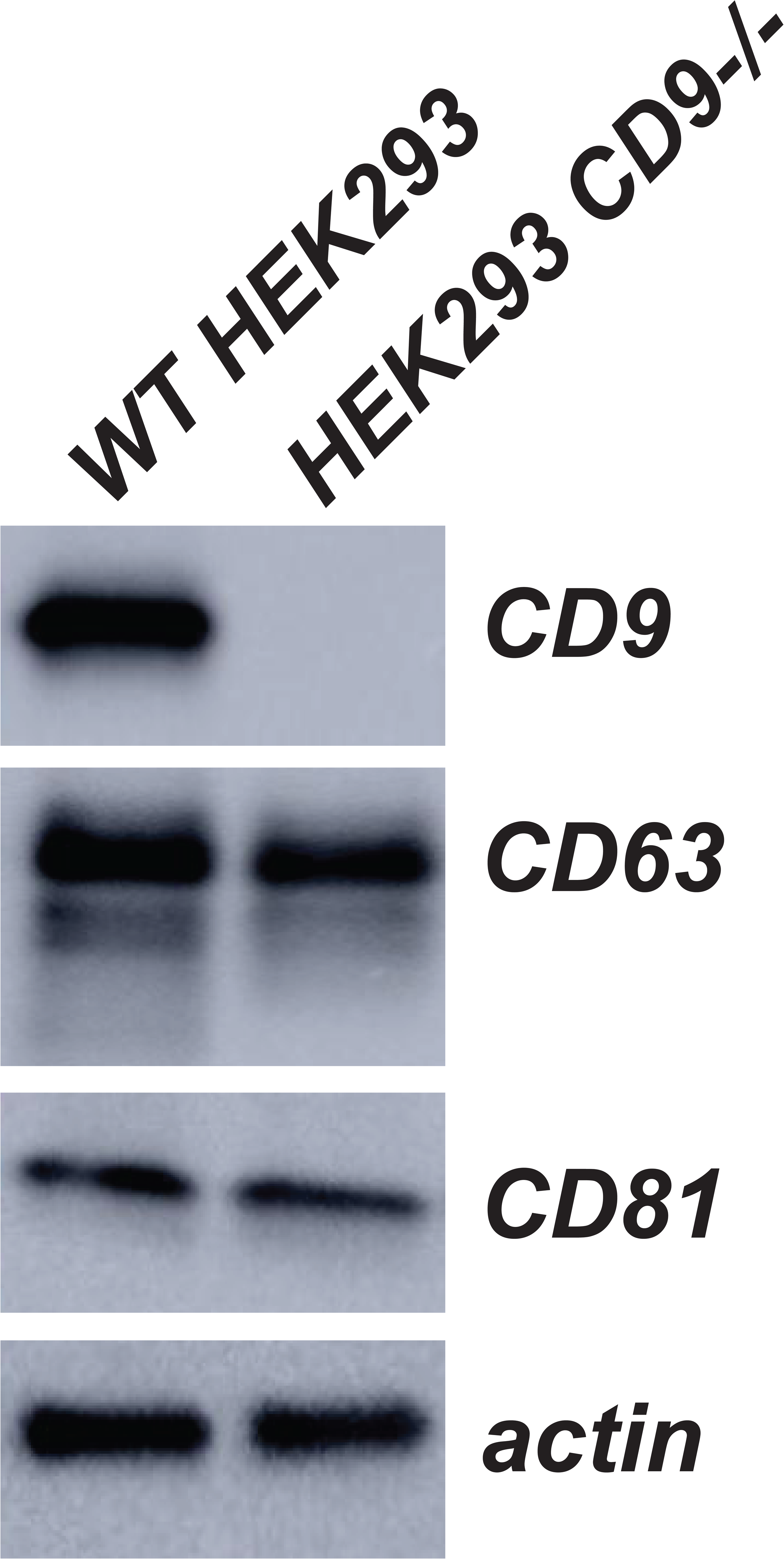
CD9^-/-^ HEK293 cells do not express CD9 protein. Immunoblot analysis of WT HEK293 cells and HEK293 CD9^-/-^ cells using antibodies specific for CD98, CD63, CD81, and actin shows that CD9^-/-^ cells do not produce CD9 protein but exhibit no apparent change in the expression of CD63, CD81, or actin.

## Abbreviations

EV: extracellular vesicles
IB: immunoblot
IFM: immunofluorescence microscopy
IPMC: intracellular plasma membrane-connected compartment
MVB: multivesicular body
SPIR: single-particle interferometric reflectance
SPIRI: single-particle interferometric reflectance imaging

